# Dynamic Behaviors of Expression Compensation between Duplicate Genes

**DOI:** 10.1101/2020.03.10.986166

**Authors:** Xun Gu

**Affiliations:** Department of Genetics, Development and Cell Biology, Iowa State University, Ames, IA 50011, USA

## Abstract

While gene or genome duplications have provided raw genetic materials for evolutionary innovations, these events have also generated massive duplicate genes, resulting in a tremendous increase to the genetic robustness of organism. Duplicate compensation indicate functional redundancies generated by gene duplications, which are widespread in all known genomes. However, the fitness trade-offs of their mutational compensation (genetic robustness) and their role in evolutionary innovation remains largely obscure. In this paper, we discuss how we can utilize the mathematical modeling approach to predicting under which condition duplicate compensation may occur. After a critical review for the models about expression dosage, compensation, and long-term survival of duplicate genes, we highlight the importance to distinguish between Function (*F*)-triggered and Expression (*E*)-triggered mechanism of duplicate compensation. Moreover, we address three fundamental questions: (*i*) Why a backup duplicate can be effectively activated by any silence mutation of the dominant duplicate, but hardly by any coding mutation resulting in impaired protein function? (*ii*) Why a dispensable duplicate gene, i.e., knockout leads to virtually no phenotype, still remains a great deal of selective constraints in the coding region? And (*iii*) under which condition expression subfunctionalization between duplicates is reversible (dosage-sharing) or irreversible (long-term survival)?

## Introduction

Functional redundancy between duplicate genes is a characteristic feature of many biological systems, ranging from cell signaling, development, to metabolism [1–16]. For instance, in signaling cascades, crucial signaling components are frequently associated with redundant, tissue-specific duplicate genes exemplified by the protein kinases, transcription regulators, as well as Wnt proteins [2, 5]. Several systematic studies in the budding yeast *S. cerevisiae* have provided comprehensive lists of functionally redundant gene pairs [17]. Gout and Lynch [13] proposed an evolutionary scenario under which the expression level of individual duplicate genes can evolve neutrally as long as they maintain a roughly constant summed expression. This mechanism allows random genetic drifts towards uneven contributions of the two duplicates to the total expression. More recently, Lan and Prichard [16] analyzed RNA-sequencing data from 46 human and 26 mouse tissues and suggested that tandem duplicates are co-regulated. Consistent with the dosage-sharing hypothesis, most young duplicates are down-regulated to match expression levels of single-copy genes. It has been that subfunctionalization of expression actually evolved not as fast as theoretically expected, which was is rare among duplicates that arose within the placental mammals [16].

Gene duplication is a fundamental process in genome evolution [1, 2, 5]. While most young duplicates became nonfunctional quickly by loss-of-function mutations (non-functionalization), mechanisms that allow some duplicate pairs to survive in the long-term evolution remain controversial [3, 8]. In addition to the well-known framework of subfunctionalization or neofunctionalization [4–9], other types of dosage constraints were also proposed [14–17]. In short, several fundamental issues have been not well addressed:

i. Why a backup duplicate can be effectively activated by any silence mutation of the dominant duplicate, but hardly by any coding mutation resulting in impaired protein function?
ii. Why a dispensable duplicate gene, i.e., knockout leads to virtually no phenotype, still remains a great deal of selective constraints in the coding region?
iii. Under which condition expression subfunctionalization between duplicates is reversible (dosage-sharing) or irreversible (long-term survival)?

The purpose of this article is to tackle these distinct yet related issues.

## Models and Methods

A class of kinetic model of gene regulation can be constructed under the conceptual framework of *E*-triggered mechanism [12], In the following we present a simple model to demonstrate some features of this mechanism, though it may not be biological very realistic.

### Before gene duplication (single-copy gene)

A simple biochemical binding reaction of the transcription factor (TF) to a given regulatory motif can be schematically represented by TF+motif ↔ TF-motif (). The kinetic model assumes (*i*) that the concentration of TF (*S*) is up-controlled by the regulatory signal, and down-regulated by the negative feed-back of the expression level; and (*ii*) that the binding-constant *z* reflects the binding (free) energy of the TF. Let *x* be the gene expression level. The regulation of gene expression can be described by the following kinetic equations, that is,

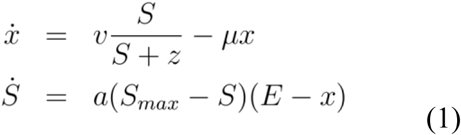

where *vS*/(*S*+*z*) is the Michaelis-Menten mechanism for the TF-motif binding; μ is the decay rate of mRNA, *a* is the increase rate of the concentration of transcription factors, denoted by [TF]; *S*_*max*_ is the maximum [TF] *in vivo*, and *E* is the optimal expression dosage.

For the long-term evolution, steady-state analysis is useful. Detailed analysis has shown that Eq.(B-1) has two stable steady states: (*i*) *x=E* and *S*<*S*_*max*_, and (*ii*) *x*<*E* and *S*=*S*_*max*_. Biologically, in the first state, the optimal expression dosage *E* is reached under the normal physiological condition, while the second state could be sub-optimal since the expression level is lower than the optimal one.

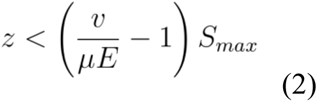

### After gene duplication

Let *x*_*A*_ and *x*_*B*_ be the expression levels of duplicates genes *A* and *B*, respectively. In the case, Eq.(B-1) can be apparently extended as follows

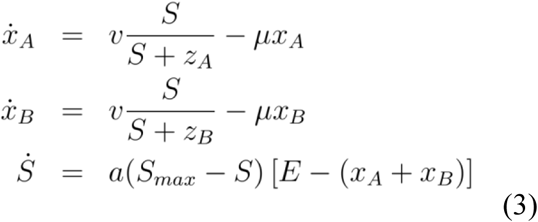

Similarly, the steady-state analysis of Eq.(B-3) has shown two stable states: (*i*) *x*_*A*_+*x*_*B*_=*E* and *S*<*S*_*max*_; (*ii*) *x*_*A*_+*x*_*B*_<*E* and *S*=*S*_*max*_. Apparently, the first state achieves the optimal expression level *E* in the duplicate pair, while the second state is sub-optimal indicating that even the expression level of two duplicates together is lower than the optimal one.

### Evolutionary model

We assume that before the gene duplication, gene regulation is under the first state so that the steady-steady expression level is *x=E*. Some studies [28–30] proposed that the binding constant (*z*) is an exponential function of mutations from the optimal binding sequence. Without loss of generality, we assume that after *t* time units since gene duplication, *z*_*A*_=*z*_*0*_ remains the optimal, whereas *z*_*B*_=*z*_*0*_ *e*^*βt*^ caused by the accumulation of mutations; the rate β represents the thermo-kinetic effect of mutations. Thus, one can show that under the constraint *x*_*A*_+*x*_*B*_=*E*, the expression ratio of *x*_*A*_ and *x*_*B*_ is given by

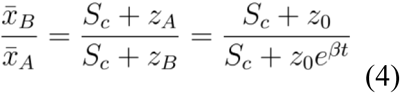

Apparently, *x*_*A*_ >>*x*_*B*_ if *z*_*A*_ <<*z*_*B*_, describing a typical case for the expression divergence between duplicates.

### Fitness effect after single-gene deletion/knockout experiment

In the case when duplicate *A* is deleted, the kinetic system of Eq.(B-3) is reduced to be the case of single-copy gene. Because of the mutational effects in the regulatory region of gene *B* that cause the increase of binding affinity (*z*_*B*_), the steady state of the expression of single gene *B* may or may not be sufficient enough to re-take the optimal dosage level *E*, that is, the first state. Thus, substituting the parameter *z* in Eq.(B-1) by *z*_*B*_=*z*_*0*_ *e*^βt^, we show that the condition is dependent of the evolutionary time *t*: For young duplicates with

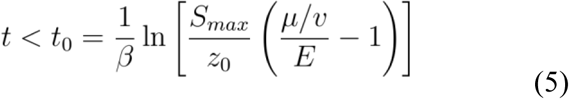

The optimal expression dosage E will be achieved. Otherwise, i.e., ancient duplicates with *t*>*t*_*0*_, the expression of gene *B* would be lower than the optimal dosage, which could affect the phenotype of the mutant.

## Results and Discussion

### Long-term duplicate compensation

Duplicate compensation indicates functional redundancies generated by gene duplications, which are widespread in all known genomes. While gene duplications result in a tremendous increase to the robustness of organisms [1–4], this type of genetic robustness may also render evolutionarily unstable [1, 5], which has only a transient lifetime because functional overlaps between duplicates would be rapidly lost, either subfunctionalization or neofunctionalization [3, 6–9]. On the other hand, numerous reports describe instances of functional overlaps between duplicates that have been conserved throughout extended evolutionary periods (see[10] for a literature survey). One such example is two duplicate O-acyl-transferases isozymes, redundantly catalyzing the conjugation of sterols to fatty acids, for which functional overlap has been conserved from yeast (Are1 and Are2) to mammals (ACAT1 and ACAT2). The challenging question is how long-term duplicate compensation can be maintained in the genome, which can be decomposed into two sub-questions: (*i*) how can two duplicate genes with redundancy be preserved against null mutations during long-term evolution, and (*ii*) how can we explain the selective constraints imposed on the coding sequences of redundant duplicates? We have realized the answer for both problems depends on the nature of signal that triggers the backup circuits between duplicate genes, which can be broadly classified two types, function (*F*)-triggered or expression (*E*)-triggered (Fig.1).

**Fig. 1.**
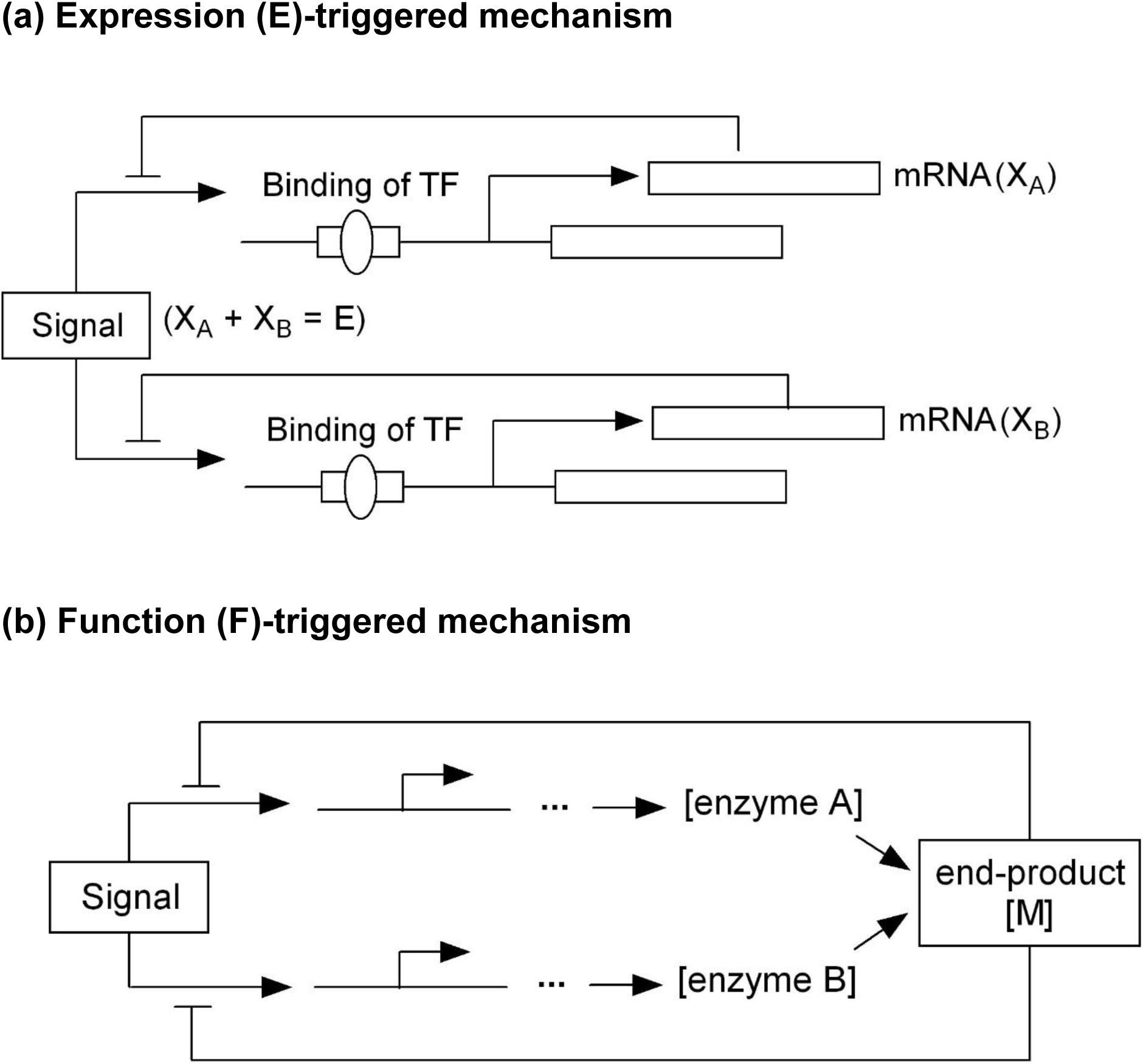
Illustration of two mechanisms of genetic robustness. (*a*) Expression (*E*)-triggered mechanism and (*b*) Function (*F*)-triggered mechanism.

### Function (*F*)-triggered backup circuit

This *F*-triggered mechanism suggests that loss of gene functionality by null mutations (either expression silencing or loss of protein function) is the signal to trigger duplicate compensation. The work by Kafri et al. [10, 11] belongs to this category.

#### Reprogramming in duplicate backup circuits

Kafri et al [11] argued that functional backup among differential expressed duplicates (*A* and *B*) may suggest that, upon null mutation in gene *A*, expression of gene *B* is reprogrammed to acquire an expression profile similar to the wild-type expression profile of gene *A*. Their model was based on the premise that the promoter architecture may be partially overlap between duplicates, as experimentally verified for the Acs1 and Acs2 isoenzymes in the yeast. Wild-type Acs1 is subject to glucose repression, but upon deletion of Acs2, the repression of Acs1 is relieved, and Acs1 acquires an Acs2-like responsiveness to glucose.

Moreover, Kafri et al [11] proposed a kinetic model, or reprogramming switch, to demonstrate the control of reprogramming process. Consider two duplicate genes (*A* and *B*) that encode enzymes *E*_*A*_ and *E*_*B*_, respectively, both of which interconvert metabolite *M*_*1*_ into metabolite *M*_*2*_, i.e., *M*_*1*_→*M*_*2*_. Though two duplicates contain the similar binding sites for a shared transcription factor (TF), only duplicate *A* is active in the wild type, because it can be induced by the TF under a low level of metabolite *M*_*1*_, whereas gene *B* can be induced only by a high level of *M*_*1*_. Upon knockout of gene *A*, metabolite *M*_*1*_ has been accumulated quickly because it cannot be efficiently interconverted to *M*_*2*_. As a result, *M*_*1*_ accumulation and the increase of TF concentration eventually result in an efficient activation of duplicate gene *B*, such that the level of enzyme *E*_*2*_ increases to interconvert *M*_*1*_→*M*_*2*_. This model provides a similar control mechanism that responses to an environmental condition (i.e., the accumulation of metabolite *M*_*1*_) or to an internal perturbation (i.e., silencing of gene *A*).

#### Responsive backup circuits (RBC) and regulatory designs

By literature surveys, Kafri et al [10] compiled a list of examples of functional redundancies between duplicate genes that have been conserved in long-term evolution. Many examples showed that a backed-up gene tends to be transcriptionally responsive to the dominant duplicate partner, and to be up-regulated if the dominant duplicate is silenced by mutations, called ‘responsive backup circuits’ (RBC). Hence, one of the two duplicates is called responsive genes because it is under repression in wild type and that repression is relieved upon its partner’s (called controller) mutation.

RBC can be illustrated by yeast Hxt gene family that encodes a set of membrane hexose transporters with different affinities toward glucose, resulting in different transport efficiencies. In yeast, glucose serves as a regulatory input for alternating between aerobic and anaerobic growth. There are two independent signaling pathways, one probing intracellular glucose concentrations and the other probing extracellular concentrations. This differential sensing shows how the responsive backup circuit of Hxt1 and Htx2 may work. Hxt2 is controlled by the feedback of two opposing signals, i.e., induction by extracellular glucose and repression by intracellular glucose. Although high glucose concentrations result in repression of Hxt2 expression, its induction could be triggered either by low environmental sugar, or alternatively, by mutations in genes responsible for glucose influx.

In short, the responsive backup circuit (RBC) theory has demonstrated the existence of a variety of kinetic mechanisms for functional backup between duplicate genes. Kafri et al [10] summarized three possible regulatory schemes. (A) Negative regulation of gene expression by its functionally redundant duplicate is a direct mechanism. (B) Substrate abundance in a pathway can activate duplicate backup gene, if the over-accumulation of substrate becomes the signal. And (C) end-product inhibition feedback inhibits both redundant enzymes for maintaining a desirable end-product level. Hence, functional loss of the dominant duplicate may result in a relief of repression for the backup duplicate.

### Expression (*E*)-triggered backup circuit

While *F*-triggered mechanism provides valuable insights about the kinetic property of duplicate compensation, we have realized that an evolutionary puzzle may emerge. As illustrated by Fig.1, the *F*-triggered backup circuits [10] can be triggered by the signal of null mutations leading to either gene silence or impaired protein function. This model predicts that functionally redundant duplicates can accumulate loss-of-function mutations in both coding and regulatory regions. Consequently, for a pair of functionally redundant duplicates that have undergone differential expressions, one of them would ultimately become a pseudogene due to the accumulation of loss-of-function mutations in the coding region. This conclusion is contradicted with the fact that many duplicate pairs remain at least partial functional overlaps for a very long span of evolutionary time. Moreover, though nonessential duplicate genes (i.e., knockout results in no apparent phenotype) are subject to high incidents of gene loss during the long-term evolution, the protein sequence conservation, as measured by the ratio of nonsynonymous rate to synonymous rate, is still usually significantly stronger than the neutral expectation.

To resolve this puzzle, Gu [12] proposed an alternative model called expression (*E*)-triggered backup circuits, suggesting that the absence of gene products (protein molecules or mRNAs) may play a key role (Fig.1). Different from the *F*-triggered backup circuits, the *E*-triggered backup circuits can only be activated by the silence of the dominant duplicate A. Since null regulatory mutations occurred in duplicate A are nearly-neutral, this duplicate gene may have a nontrivial chance to be lost during evolution. On the other hand, when gene *A* encodes a functionally null protein, the accumulation of mutated mRNA can effectively prevent the backup activation. Consequently, deleterious mutations in the protein sequence must be eliminated by the purifying selection. Therefore, *E*-triggered backup circuits explains protein sequence conservation in those genes with high incidence of gene gain and loss across species.

### Kinetically reversible expression compensation between duplicates

The e*EK* (expression-triggered evolutionary kinetics) represents a class of evolutionary models that highlight the underlying kinetic processes of molecular interactions (BOX-1).

#### Evolutionary steps from reversible to irreversible subfunctionalization

From the systems biology view [18–21], modeling the *E*-triggered mechanism should involve the kinetics of expression that may differ between duplicates. Based on the mathematical models of gene regulation [22–24], the duplicate copy with low expression can be kinetically inhibited by the other copy with high expression. When the highly-expressed duplicate copy is deleted, the expression of lowly-expressed copy can be kinetically elevated for compensation, a phenomenon called *reversible subfunctionization*. For two duplicate genes that have diverged *t* evolutionary time units ago, a complete *eEK* analysis in BOX-1 yields the following evolutionary scenarios (Fig.2).

- *Initial stage:* Let *E* be the ancestral expression level of the single progenitor gene under a certain tissue/developmental stage. After the gene duplication, the two identical copies have the same expression level *E*/2, respectively.
- *Early stage:* The early stage can be defined by the time *t*<*t*_*0*_, where the criteria *t*_*0*_ is determined by the *EK* model. During this stage, reversible subfunctionalization may occur. Suppose that the transcription of duplicate gene *B* becomes kinetically less efficient in a given tissue, whereas duplicate *A* is dominant in expression. For young duplicates with the age *t* < *t*_*0*_, when duplicate *A* is silenced, the expression level of duplicate *B* would be kinetically forced back to the normal level (*E*). In other words, the expression subfunctionalization is almost completely reversible.
- *Later stage* (*t* > *t*_*0*_): Since the expression of gene *B* is trivial in the given tissue, the transcription efficiency of gene *B* is decreasing with the evolutionary time *t* by the accumulation of regulatory mutations. Consequently, when *t*>*t*_*0*_, the expression of gene *B* is no longer able to reach the optimal level *E* when duplicate *A* is silenced. Hence, the subfunctionalization becomes partially reversible.
- *Final stage:* After the long evolutionary time period (*t* >> *t*_*0*_), duplicate *B* becomes totally inactive under this physiological condition, leading to a complete loss of functional compensation, or irreversible subfunctionalization.

**Fig. 2.**
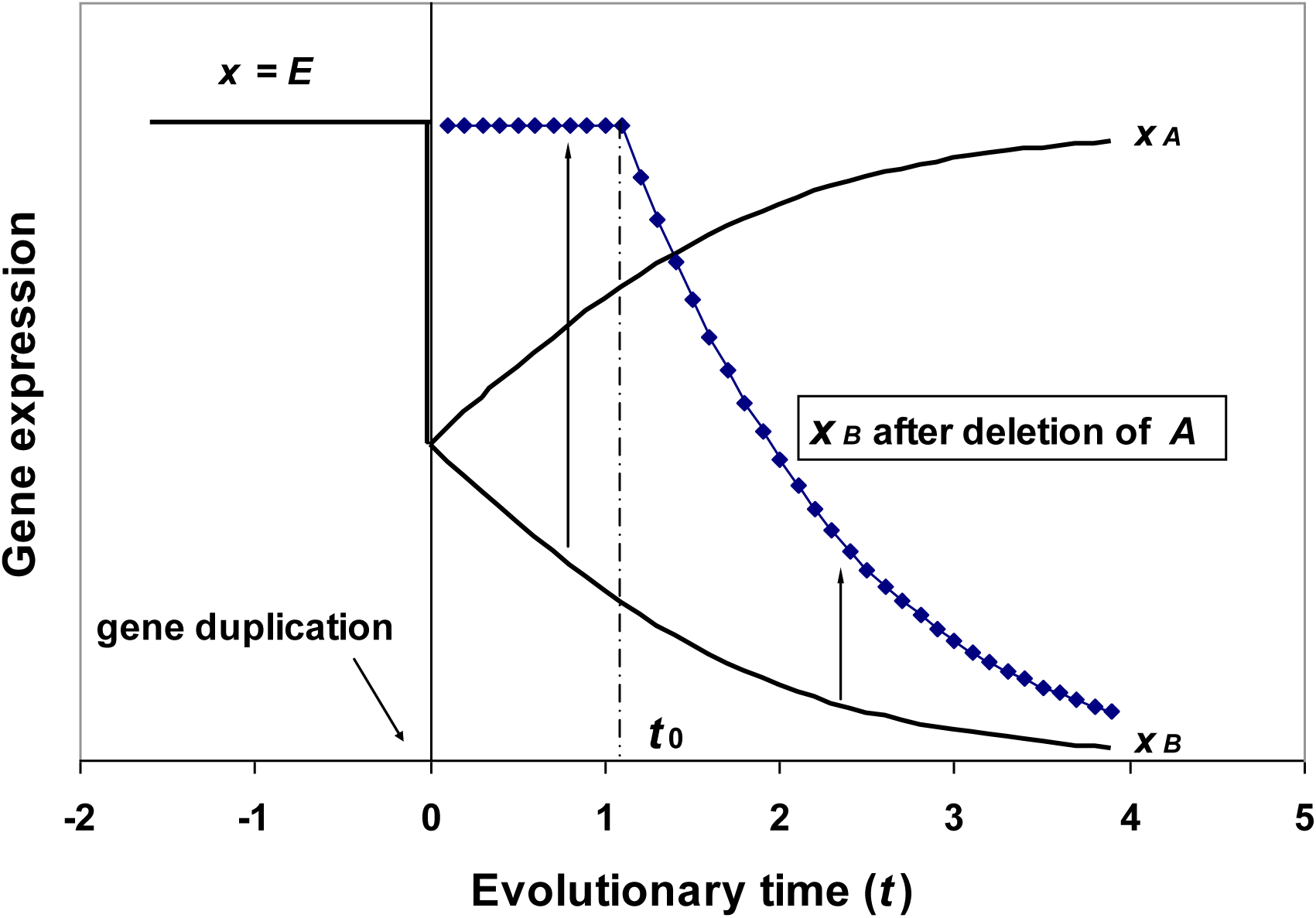
The E-triggered evolutionary kinetics (e*EK*) model between duplicates. Consider a given condition (cell-type, tissue or development). Initially, the gene expression is at the optimal dosage *E*. After gene expression, the expression of gene *A* becomes dominant, while that of *B* becomes minor due to the accumulation of mutations in the regulatory region, or epigenetic factors. During the course of evolution, the regulatory feedback reveals two distinct kinetic properties when gene *A* is silenced: for young duplicates (*t* < *t*_*0*_), the expression of gene *B* would be elevated to the normal dosage *E*, or called reversible subfunctionization; otherwise (*t* > *t*_*0*_), it only achieves a sub-optimal dosage that may cause fitness reduction.

#### Rapid divergence in the expression profile leads to reversible subfunctionalization

In short, reversible subfunctionalization can emerge rapidly. Under the normal condition, the designed expression level of the target gene has been set by the up-level regulatory network. After gene duplication, both target duplicate and backup duplicate have the kinetic capacity to reach the designed expression level, which can evolve quickly toward differential expressions. That is, while the target duplicate is responsible for to the designed level, the backup duplicate keeps a low expression level, as a result of kinetic competition.

### E-triggered mechanism prevents deleterious mutations in protein function

Suppose that two young duplicated genes *A* and *B* with reversible subfunctionalization in their expression patterns of tissues *a* and *b*, respectively. That is, silence of gene *B* would kinetically turn on the expression of gene *A* in tissue *b*, and *vice versa.* Consider the case when a loss-of-function mutation occurred in protein *B* (marked by *), leading to the expression of nonfunctional protein *B*\*. Under the *EK* model, the reversible process for expressional compensation is triggered by no expression of the gene, rather than the impaired protein function. As a result, gene *B*\* (with a loss-of-function mutant in the coding region) cannot be functionally compensated by gene A. Because expression of nonfunctional protein *B*\* in tissue *b* may have deleterious phenotypes, this mutant has to be removed from the population by the purifying selection.

### A speculation on a general E-triggered model

The E-triggered model for genetic robustness has reasonably explained why a backup duplicate can be triggered only by the silence mutation, why dispensable gene remains sequence conservation in the coding region, and why subfunctionalization in expression is reversible between young duplicates. However, the major challenge this model is facing is how to identify the key-step for triggering among multiple cellular processes from the initiative of transcription to the final product (protein) at the targeted subcellular location. Tentatively, we speculate that it might be related to another important mechanism of genetic robustness, the molecular chaperon-mediated buffering [23, 25].

Molecular chaperones, such as Hsp70, are proteins that assist the covalent folding and the assembly of other macromolecular structures, which plays a significant role to increase organismal robustness [23]. Ample evidence has demonstrated that molecular chaperons can buffer the phenotypic effects of mutations by mediating folding of mutant proteins (misfolding avoiding), alleviating the deleterious effects of mutations. In sufficient activity of molecular chaperones may inflict a deregulation of cellular signaling systems [25]. Moreover, molecular chaperons may facilitate the structural evolution of proteins. By maintaining nonnative proteins in a soluble, folding-competent state, chaperones may broaden the range of mutant proteins subject to positive selection [26, 27]. On the other hand, molecular chaperones can sensitize highly unstable proteins and direct them towards degradation instead of folding, avoiding the accumulation of severe deleterious mutations.

We suspect that molecular chaperons may also sensitize the absence of target proteins (Fig.3). For instance, the concentration of chaperon-protein complexes may be the sensor of *E*-trigger mechanism of genetic robustness. In the case of gene silence, it is virtually zero, forcing an increase of the up-level regulation signal (*S*) to trigger backup pathways, as shown in BOX-1 in the case of duplicate compensation. We acknowledge that the model is subject to some speculations, which will be further explored in our future study.

**Fig. 3.**
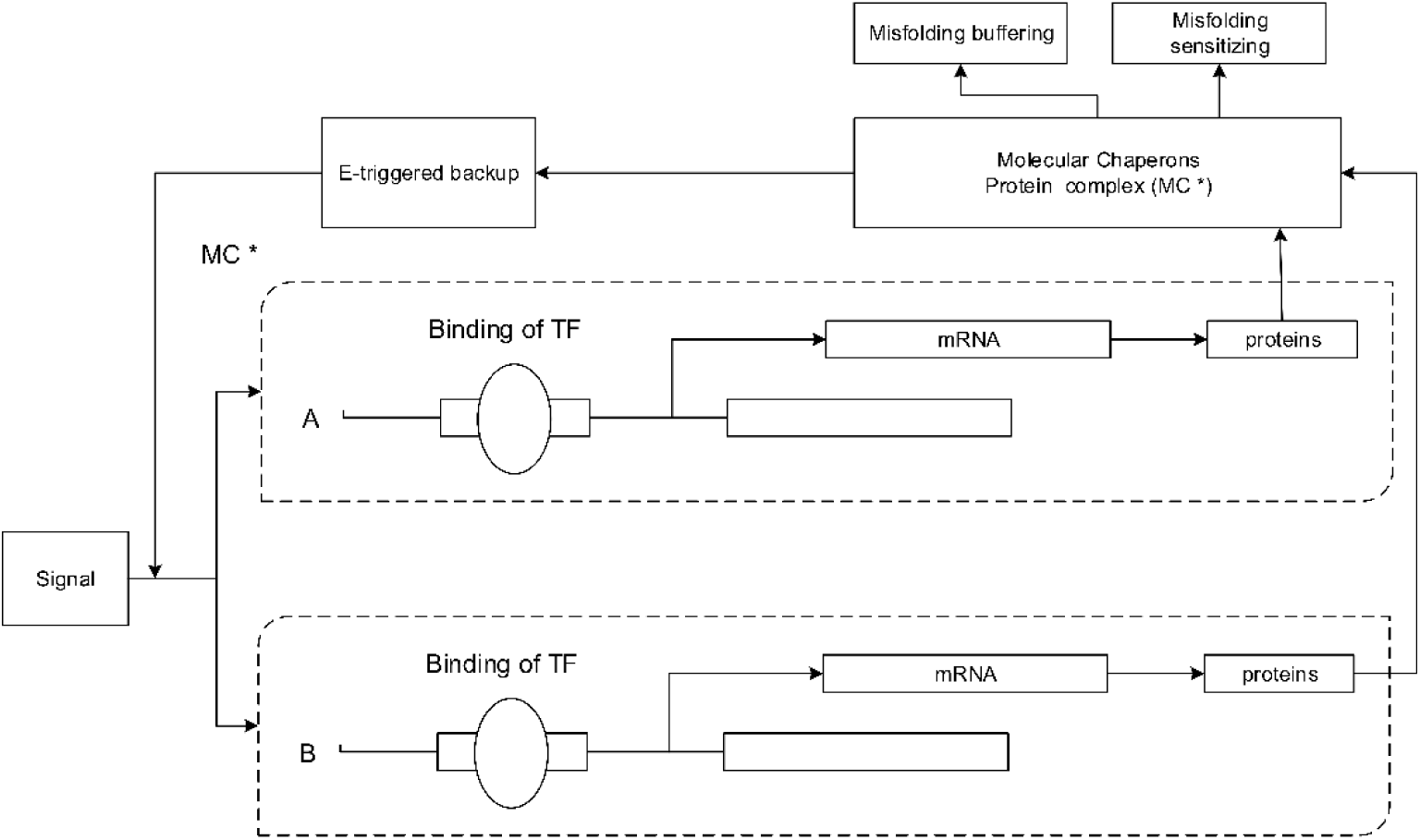
The molecular-chaperon model for E-triggered mechanism of genetic mechanism. Molecular chaperons may also sensitize the absence of target proteins through instance, the concentration ratio of the released (well-folded) proteins to chaperon-protein complexes may be the sensor of *E*-trigger mechanism of genetic robustness. In the case of gene silence, this ratio is virtually zero, forcing an increase of the up-level regulation signal (*S*) to trigger backup pathways,

## Acknowledgements

This work was partly supported by the fund from Iowa State University.

